# The effect of EEG lead configuration on early TMS–EEG artifacts

**DOI:** 10.64898/2026.02.10.705170

**Authors:** K Lankinen, G Fadel, A Nummenmaa, RJ Ilmoniemi, T Raij

## Abstract

Transcranial magnetic stimulation (TMS) combined with electroencephalography (EEG) has strong potential for recording cortical reactivity and connectivity. However, this promise is hampered by TMS-induced EEG artifacts. Here, we examine the origins of these artifacts with phantom TMS–EEG recordings and simulations. We focus on two major types of artifacts: (1) the TMS pulse artifact during each ∼0.2 ms TMS pulse and (2) the decay artifact that may last tens of milliseconds. We examine how these artifacts change as a function of the relative position between TMS coil windings and EEG electrode leads. We also examine the hypothesis that certain EEG lead configurations may reduce or even cancel out these artifacts. In experimental results across 23 different TMS coil / EEG lead configurations, the amplitudes between the TMS pulse artifact and the decay artifact were highly correlated (Spearman ρ = 0.86, *p* < 0.001), suggesting that the decay artifact is caused by the TMS pulse artifact. As predicted, in certain EEG lead configurations, both the TMS pulse and decay artifacts were minimized. The simulations confirmed that the TMS pulse artifacts depended on the electromagnetic induction from the TMS coil windings to the EEG leads. These results illuminate the generator mechanisms of—and possible means to reduce—both artifacts.

## Introduction

Transcranial magnetic stimulation (TMS) in combination with electroencephalography (EEG) provides a way to measure cortical reactivity and connectivity with high temporal resolution (Ilmoniemi et al., 1999). A practical major issue hampering this approach is the TMS artifacts in the EEG signal (Virtanen et al., 1999; Ilmoniemi and Kicić, 2010; Rogasch et al., 2013; Ilmoniemi et al., 2015; Varone et al., 2021; Hernandez Pavon et al., 2023; Stango et al., 2025). In this study, we focus on two TMS–EEG artifacts: the *TMS pulse artifact* and the *decay artifact*. The third major type of TMS–EEG artifacts, the scalp muscle artifact, is outside the scope of this study.

First, each TMS pulse, typically lasting ∼0.2 ms, creates an early and very short *TMS pulse artifact* during the pulse. It likely results from direct electromagnetic induction by the TMS pulse into the EEG electrodes and leads (Ilmoniemi et al., 2015). Ideally, this artifact recorded with EEG would have the same ∼0.2 ms duration and waveform (*e.g*., biphasic) as the TMS pulse itself. In practice, in EEG recordings this artifact appears temporally slightly stretched (up to about 0.55-ms duration), depending on the EEG low-pass filter settings allowed by the sampling rate and amplifier design. This artifact is also very strong, as it may reach peak amplitudes of several volts, i.e., orders of magnitude larger than the microvolt-range EEG signals from the brain. None of the current commercially available TMS–EEG systems can capture this artifact as faithfully as MHz-sampling oscilloscopes, but with suitable settings (sampling frequency ≥ 20 kHz, lowest low-pass filter cutoff frequency ≥ 3 k Hz, DC coupling), some state-of-the-art EEG instruments can record it reasonably well (Jamil et al., 2024).

Second, there is the *decay artifact* (Ilmoniemi et al., 2015; Stango et al., 2025) that can last tens of milliseconds after the TMS pulse. This exponentially decaying signal likely has a more complex, largely capacitive, origin, where the TMS pulse charges several parts of a distributed patient – electrode – EEG lead – EEG amplifier system; this energy discharges relatively slowly towards the patient (via the scalp-electrode interface through conducting paste) and in the EEG amplifier (with high-impedance input stage and internal RC circuits). The decay artifact is much weaker than the TMS pulse artifact but still orders of magnitude stronger than endogenous brain activity measured with scalp EEG.

Even with the best currently available EEG technologies and techniques, both TMS pulse and decay artifacts are present in TMS–EEG recordings (Stango et al., 2025). At typical TMS intensities (∼100% resting motor threshold, rMT) they dominate over EEG signals from the brain in the time window where the physiologically most interesting events occur. First, reactivity is best quantified from the initial neuronal activation to the TMS pulse, observed in EEG within ∼3 ms after the pulse (Beck et al., 2024). Second, connectivity is best measured by quantifying the conduction delays from the initial activation site to secondary activation locations outside the primary target (Cracco et al., 1989; Ilmoniemi et al., 1997; Hernandez-Pavon et al., 2023). While most conduction delays in humans are unknown, they typically fall in the 4 – 20 ms range, which overlaps with the decay artifacts. One could argue that the TMS pulse artifact is less interesting because it ends before neuronal activations begin. However, since the TMS pulse artifact kicks off the sequence of later and longer-lasting artifacts that do matter, understanding it may offer clues for minimizing all types of TMS-related artifacts. Previous human studies have suggested that reducing the induction from the TMS coil to the EEG leads by carefully orienting the EEG leads relative to the TMS coil windings may reduce the decay artifact amplitude and duration (Sekiguchi et al., 2011; Hernandez-Pavon et al., 2023). However, quantification with simulations and phantom recordings while parametrically varying the relevant variables has not been done previously. Moreover, the relationship between the TMS pulse artifact and the decay artifact has not been quantified before. Therefore, we here report on a single-channel EEG experiment where we systematically vary the TMS coil position and EEG lead configuration in a spherical phantom, using both recordings and simulations. This illuminates the relationship between these factors, which may guide future expansion to whole-head multi-channel EEG systems in humans.

## Methods

### Phantom

We used a spherical MRI phantom (radius *R*_s_ = 9.1 cm) made of a non-conducting plastic enclosure filled with liquid. For the experiment, the phantom was covered by a cotton fabric cap soaked in a 0.9% saline solution (**Figure 1**). EEG electrodes were attached on the cap using conducting paste (Grass EC2, Natus Medical Incorporated, Middleton, WI, USA). Next, the phantom was tightly wrapped in non-conducting plastic film to prevent drying of the cap and the EEG paste and to keep the electrodes in place. Finally, a TMS navigation tracker was attached using a 3D-printed custom adaptor piece between the tracker base and phantom. **Figure 1** shows the phantom setup.

**Figure 1.**
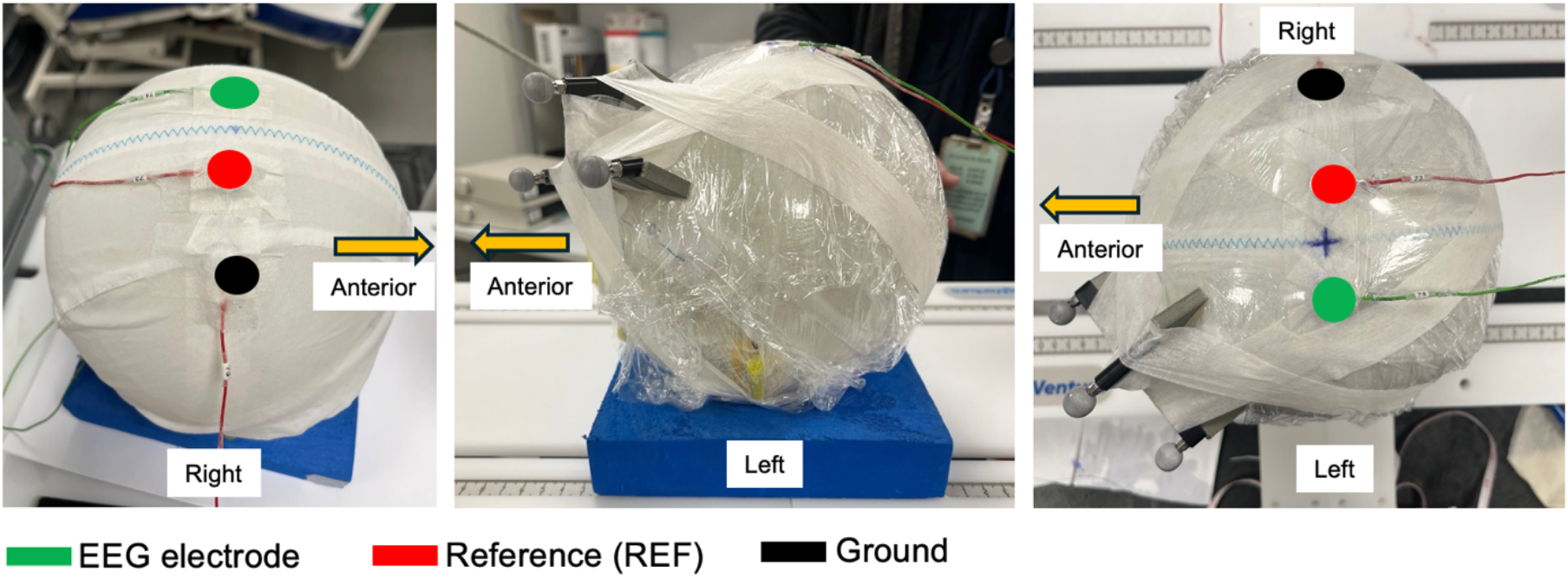
A spherical TMS–EEG phantom. **Left:** A view from the right upper angle. The phantom was covered with a cotton cap that was soaked in 0.9% saline solution. Three electrodes were attached on the cap: EEG, reference (REF), and ground (GND). **Middle:** A view from the left. After wrapping the phantom in plastic film, a TMS navigation head tracker was taped on the surface. **Right:** A view from top.

### Electrode setup

One EEG electrode, reference (REF), and ground (GND) were placed on the spherical surface so that the midsagittal line of the sphere was halfway between EEG and REF electrodes (**Figure 2**). The distance between EEG and REF was *d* = 40 mm along the spherical surface. The coordinate system defined the origin (0,0,0) at the center of the sphere. The TMS coil was placed above the phantom horizontally, slightly above the electrodes to prevent mechanical vibration artifacts from the coil. Specifically, distance from the coil housing outer surface to EEG electrodes was 5 mm (from coil winding midpoint to EEG electrodes 10 mm).

**Figure 2.**
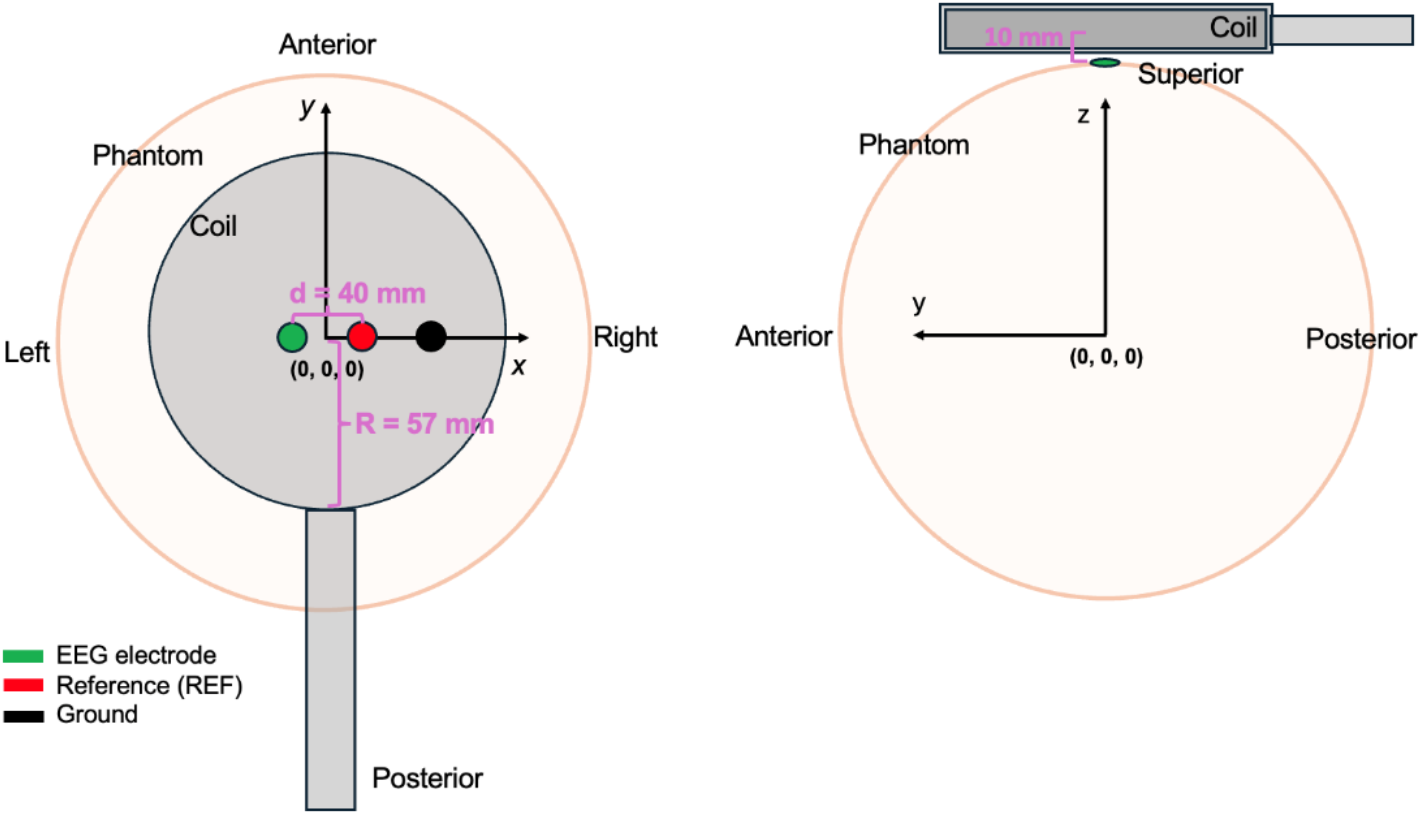
Schematics and coordinate system of the experiment. The radius of the sphere was *R*_s_ = 91 mm and the radius of the coil outer windings *R* = 57 mm. The EEG and the REF electrodes were placed symmetrically across the midsagittal line with a distance *d* = 40 mm between them. GND was placed 40 mm right to the REF.

### EEG lead configurations and TMS coil positions

We measured single-pulse TMS (spTMS) evoked EEG responses in 23 different EEG lead configurations and TMS coil positions (**Figure 3**). The EEG and REF leads were either non-crossed (Conditions 1–12) or crossed (Conditions 13–23). For both non-crossed and crossed lead configurations, the EEG and REF leads eventually met and were thereafter twisted in a pair until connecting to the EEG jackbox. This setup formed lead “loops” from the electrodes to the start of the twisted pair. The length of the loop *h* measured on the *y*-axis from the electrodes to the point where the twisted pair started, was parametrically varied as multiples of the coil radius *R* (*h* = 0.5, 1, 1.5, or 2*R*). The only exception was that for crossed Conditions, *h* = 1*R* was in practice not possible due to inflexible wire segments close to the electrodes. The TMS coil positions were varied in 3 steps along the *y*-axis. Specifically, the coil center was positioned on top of the electrodes (center), or with the electrodes on the lower or upper edge of the coil windings. In the crossed set (Conditions 15, 19, and 23) a 4^th^ condition with the wire crossing at the TMS coil center was included.

**Figure 3.**
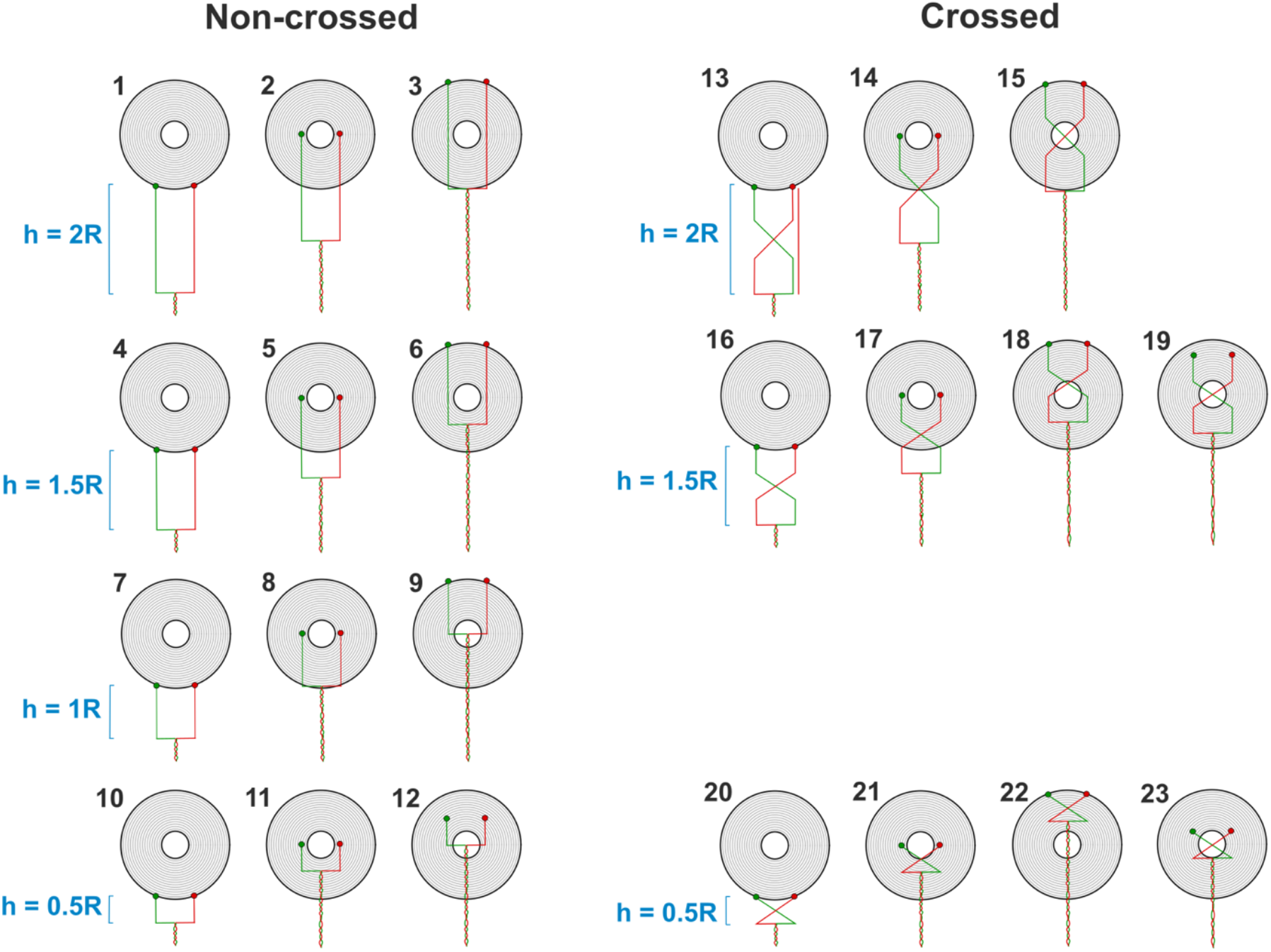
EEG wire configurations and TMS coil positions. The leads were non-crossed (Conditions 1–12) or crossed (Conditions 13–23). The EEG lead loop length *h*, from the electrodes to the start of the twisted pair, were varied as *h* = 0.5, 1, 1.5, and 2 times of the coil radius *R*.

### TMS, MRI, and navigation

TMS was delivered with a MagPro X100 stimulator and circular MC-125 coil (MagVenture, Farum, Denmark). The pulse strength was 76 A/μs (50% of the maximum stimulator output). The waveform was biphasic with pulse duration of 0.28 ms and recharge delay of 150 ms. For each condition, 10 single pulses were delivered with one pulse every 2 seconds.

The TMS coil location relative to the phantom and EEG electrodes/leads was planned and recorded with a TMS neuronavigation system (LOCALITE GmbH, Bonn, Germany) with an optical camera and passive trackers (Polaris Spectra, Northern Digital Inc., Waterloo, Ontario). For the phantom, T1-weighted anatomical images with 3 navigation capsules were acquired with a multi-echo MPRAGE pulse sequence (TR = 2510 ms; 4 echoes with TEs = 1.64 ms, 3.5 ms, 5.36 ms, and 7.22 ms; 176 sagittal slices with 1 × 1 × 1 mm^3^ voxels, 256 × 256 mm^2^ matrix; flip angle = 7°) in a 3 T Siemens Trio MRI scanner (Siemens Medical Systems, Erlangen, Germany) using a 32-channel head coil.

### EEG recordings and analysis

EEG was recorded with a Bittium NeurOne Tesla EEG system (Kuopio, Finland) in DC coupling mode, digital lowpass filter cutoff frequency of 5 kHz, anti-aliasing analog lowpass filter cutoff frequency of 3.5 kHz, and sampling rate of 20 kHz. The 10 individual responses for each condition were averaged within Conditions.

To quantify the strength of the artifacts, we calculated the area under curve (AUC) for both TMS pulse and decay artifacts (**Fig. 4**). Because the artifacts had both positive and negative values, we calculated the AUC value for the absolute values of the signals. In addition, we quantified the maximum amplitude of the TMS pulse artifact.

**Figure 4.**
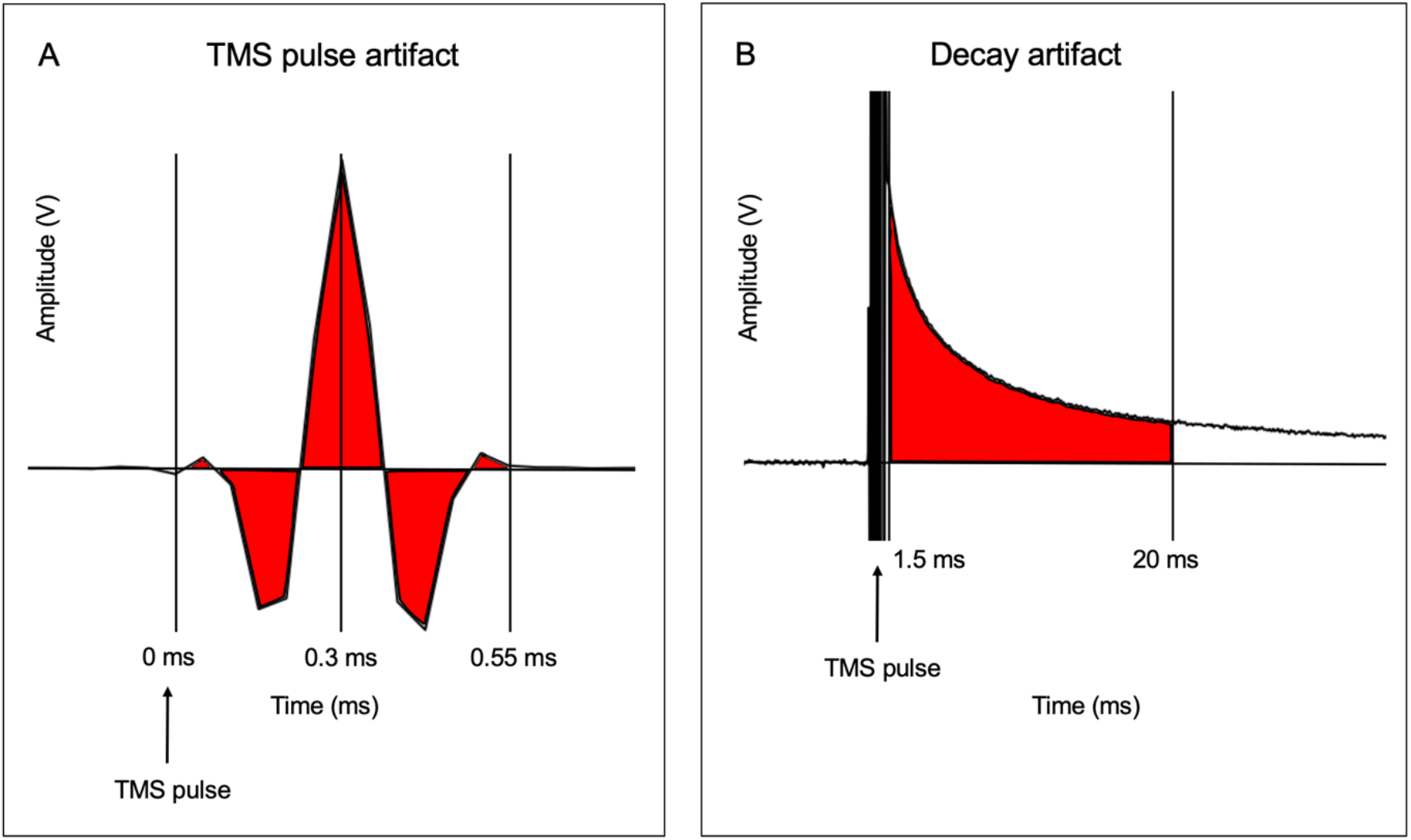
Quantification of TMS pulse and decay artifacts. We calculated the area under curve (red) A. between 0–0.55 ms for TMS pulse artifacts and B. 1.5–20 ms for decay artifacts. In addition, we quantified the peak of the TMS pulse artifact at 0.3 ms (A).

**Figure 5.**
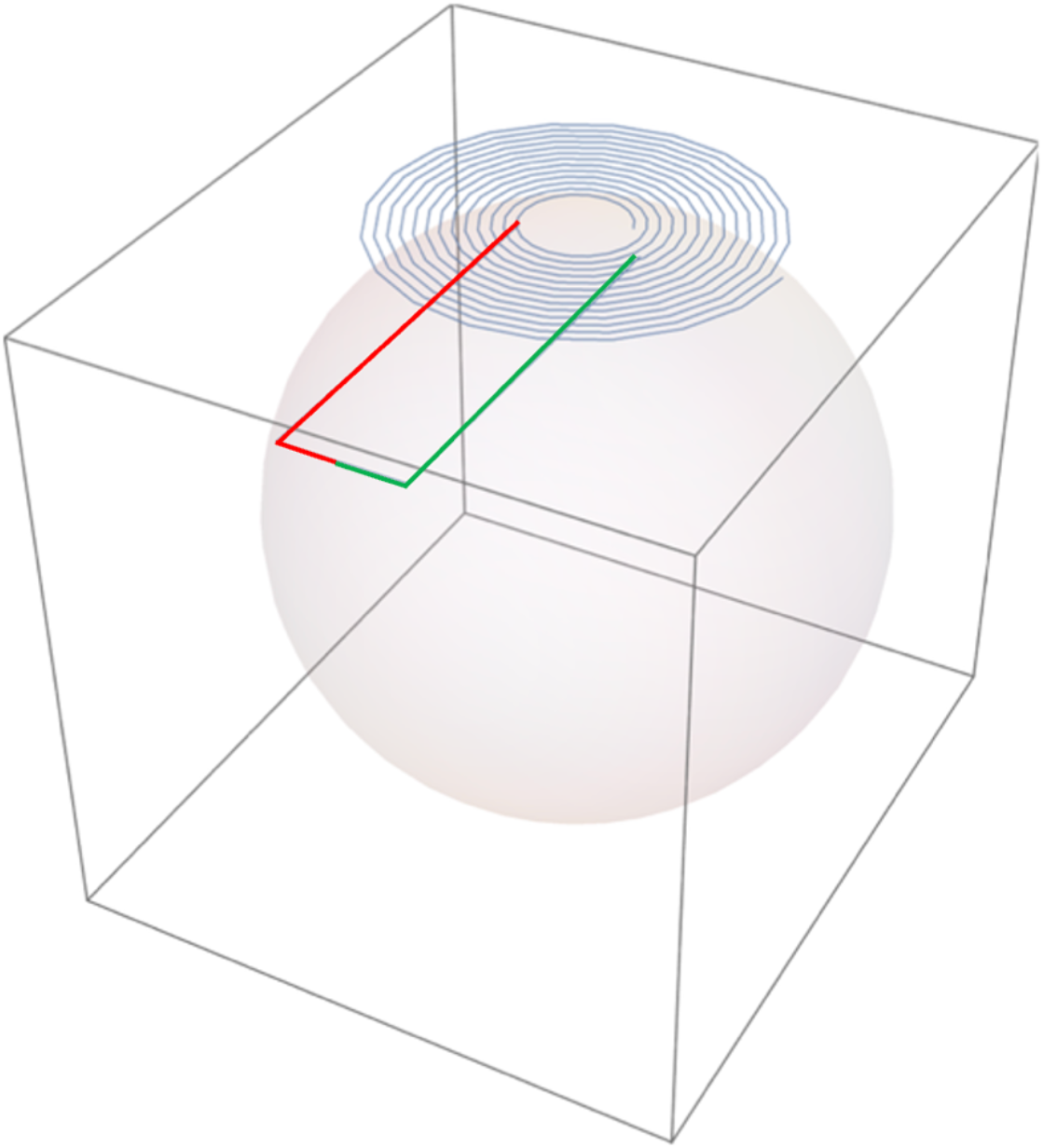
Simulation of the EEG response caused by TMS pulse, illustration of the setup. The coil is placed over the sphere in the *x–y* plane. The EEG leads are here according to Condition 1 (Figure 3).

### Simulations

We additionally simulated the amplitude of the TMS pulse artifact in the EEG leads using the same lead configurations and coil positions as in the recorded data (Fig. 3). The real TMS navigator data were used to confirm the TMS coil locations relative to the phantom. In this model, the amplitude of the pulse artifact in the EEG signal depended on the phantom radius (and thus the locations of the electrodes with respect to sphere center), lead configuration, coil position and orientation (always in the *x*–*y* plane), and TMS pulse intensity, with the artifact voltage being proportional to the time derivative of the coil current, i.e., *dI*/*dt*. The phantom was a spherically symmetric two-layer homogeneous isotropic volume conductor (mimicking scalp and skull) with a conductivity of 0.33 S/m in a 1-mm-thick surface layer and 0 S/m in the volume inside this layer. Because of the spherical symmetry, the spherical model was used for calculating the induced voltages between the two electrodes. Since the exact conductivities and layer thicknesses do not influence the induced voltages, these parameters were not explicitly considered in the calculations. However, the relative locations of the center of the sphere, the electrodes and coil must be taken into account. The EEG amplifier measures the voltage induced in the loop defined by the electrode leads and the path of current in the conducting sphere from one electrode to the other. One can intuitively visualize the current path in the phantom between the electrodes by imagining that current is fed into the conductor via the electrodes; the path is then a continuous distribution of current. The key simplification is that, because radially oriented primary currents in the spherical model do not produce an external magnetic field, the three-dimensional current path can be replaced by two line segments of virtual lead: one from the first electrode to the center of the sphere and the other from the center of the sphere to the second electrode. The induced voltage is then *dI/dt* times the mutual inductance between the TMS coil windings and the loop defined by the electrode leads and these two current segments.

## Results

Figure 6 shows the recorded TMS pulse and decay artifacts for all Conditions. The TMS pulse artifact was observed at 0–0.55 ms, with amplitudes (max ∼0.45 V) and polarities varying across the Conditions. The decay artifacts were maximal during the first 20 ms (but continued tens of ms longer), again with the amplitudes varying across Conditions. 14 out of 23 Conditions showed negative decay artifact values.

**Figure 6.**
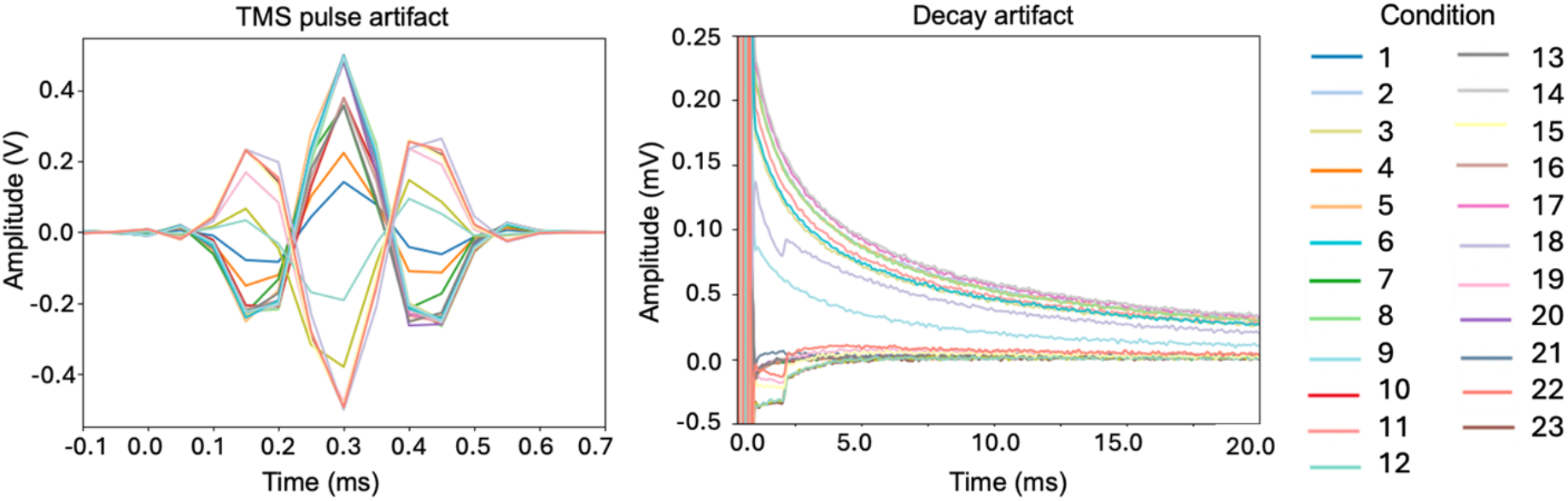
Recorded TMS pulse and decay artifacts for the 23 di[erent Conditions.

Figure 7 shows quantification of the TMS pulse artifacts (magenta) and decay artifacts (black). The figure shows that for most of the Conditions, the TMS pulse and decay artifact amplitudes were correlated. The biggest difference between the artifacts was in the crossed Conditions that had the smallest wire loop (*i.e*., Conditions 21, 21, 23, 24). Across all Conditions, the TMS pulse and decay artifacts were correlated (Spearman ρ= 0.86, *p* < 0.001).

**Figure 7.**
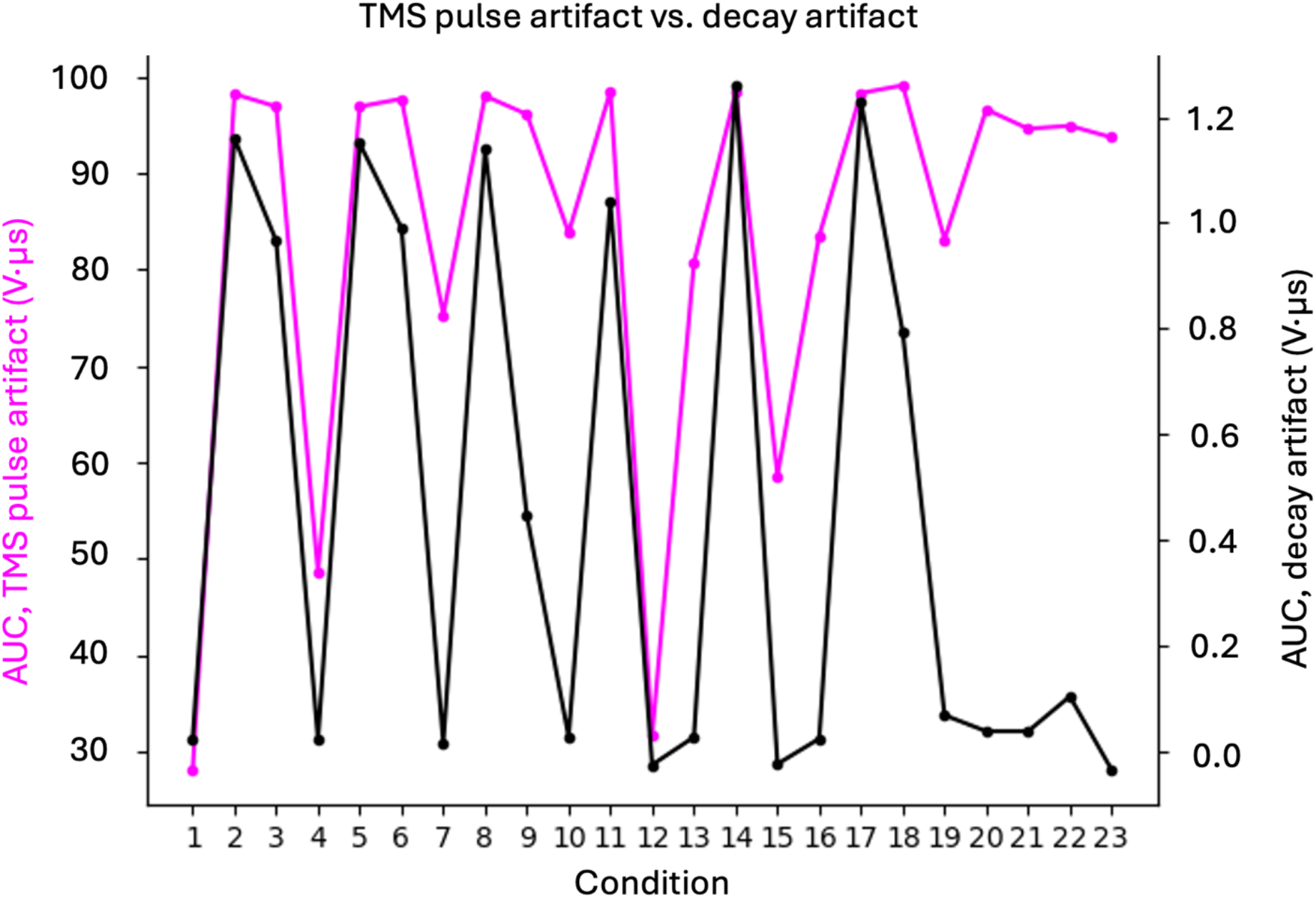
Recorded data, amplitude comparison between the TMS pulse artifacts and decay artifacts. The amplitudes were quantified as AUC values for TMS pulse (magenta) and decay (black) artifacts as shown in **Fig. 4**.

Figure 8. shows quantification of the recorded TMS pulse (peak value at 0.3 ms) artifacts (green) and simulations (orange). The figure shows that for most of the Conditions, the artifacts were correlated. Across all Conditions, the TMS pulse and simulations were correlated (Spearman ρ = 0.90, *p* < 0.001).

**Figure 8.**
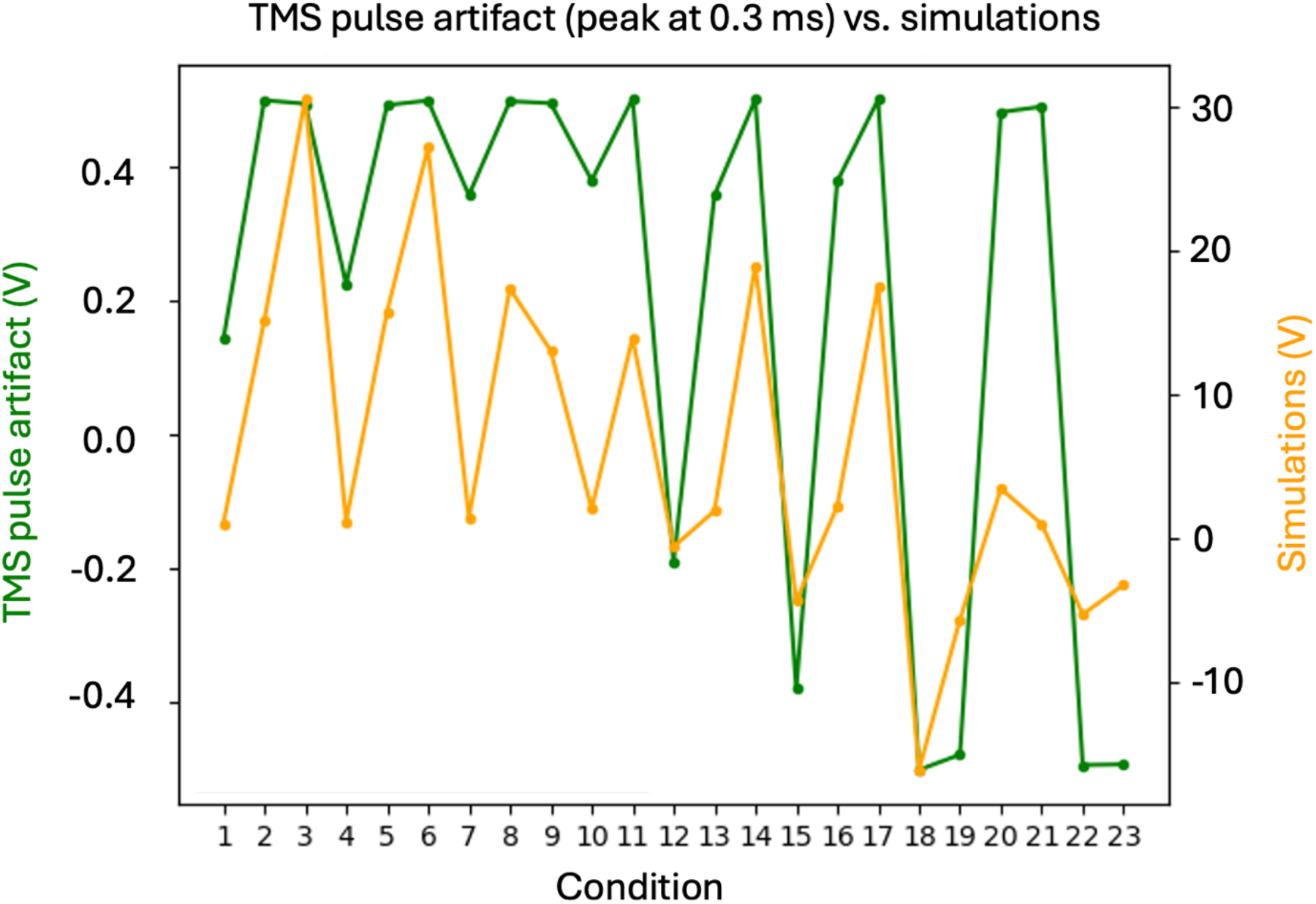
Recorded versus simulated data amplitudes for the TMS pulse artifacts. The simulated values very closely corresponded to the recorded data, reflecting that the TMS pulse artifact results from direct electromagnetic induction from the TMS coil to the EEG electrodes and leads.

Figure 9. shows quantification of the decay artifacts (black) and simulations (orange). The figure shows that for most of the Conditions, the artifacts were correlated. Across all Conditions, the decay artifact and simulations were correlated (Spearman ρ = 0.66, *p* < 0.001).

**Figure 9.**
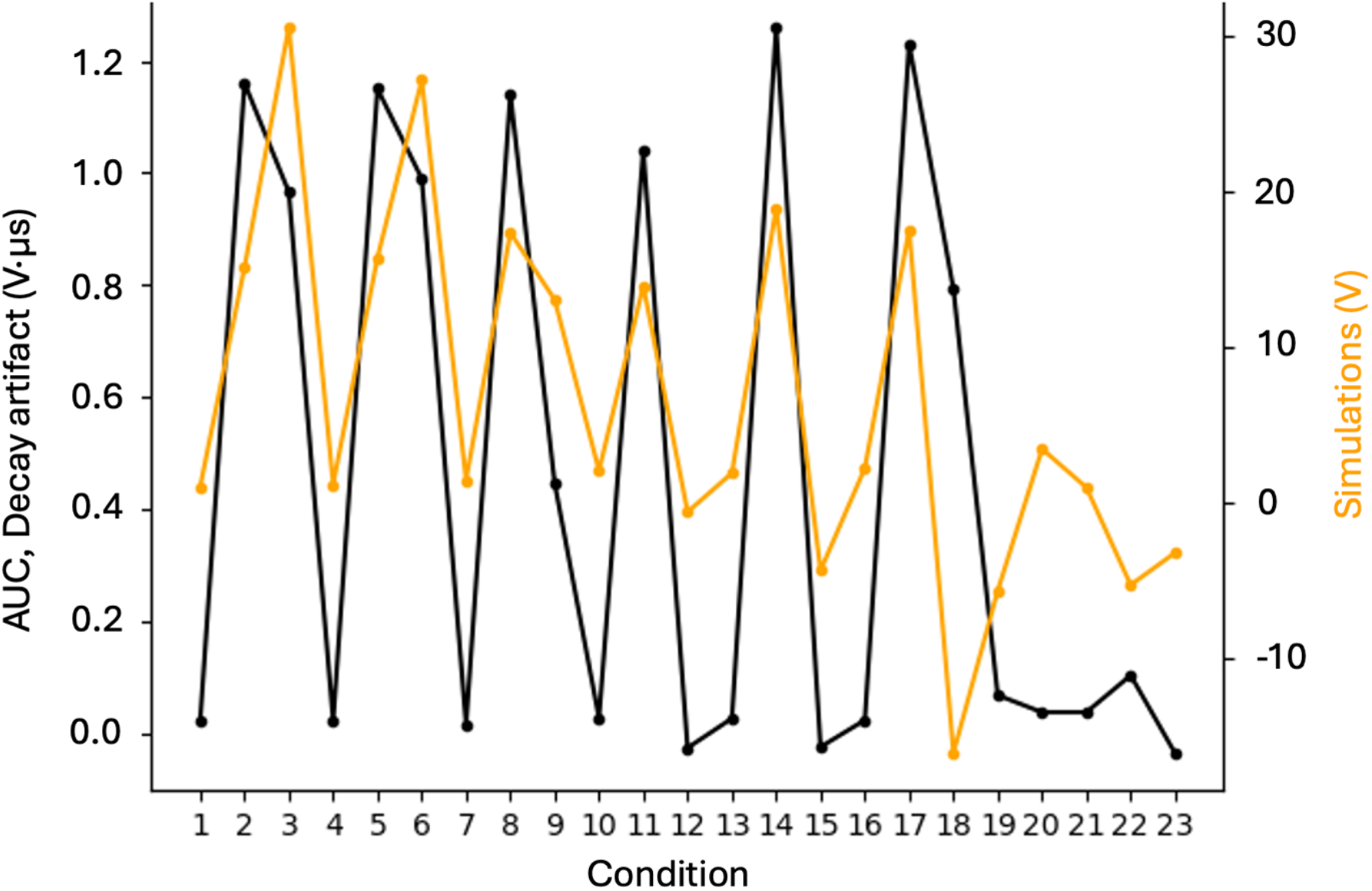
Recorded versus simulated data amplitudes for the decay artifact (black) and simulated TMS pulse artifacts (orange). The amplitudes for the decay artifacts were quantified as AUC values as shown in **Fig. 4**.

## Discussion

We systematically investigated the dependence of the TMS pulse artifact and decay artifacts on the lead configuration and coil position and compared the artifact amplitude with simulations. First, the TMS pulse artifact and decay artifact strengths were correlated, suggesting that the decay artifact is caused by the TMS pulse artifact, *i.e*., the induced voltage and consequent (small) charge redistribution across the electrodes. Second, the experimentally recorded values agreed with the simulations. Finally, we found that with certain EEG lead configurations, both types of artifacts were minimized.

The high correlation between the TMS pulse artifact and decay artifact suggests that they are closely related. The voltage pulse (which we measure as the TMS pulse artefact) charges the electrode contact (tissue–electrode–paste–metal interface), which then discharges with some time constants. However, this voltage-pulse effect is not always the same, because the electrode–skin connection can change by, e.g., drying paste, or slight movement of the electrodes. Thus, perfect correlation cannot be expected.

The simulations and recorded values correlated quite strongly, which helps us understand the results in terms of variables involved in the simulation. However, theory simplifies many things, such as the effect of electrode contacts. In addition, there is inevitably some inaccuracy, which arises, *e.g*., from setting up the wires and electrodes and positioning the coil. Thus, we cannot expect a perfect correlation between simulations and recordings either.

The promise of using TMS–EEG for quantifying cortical reactivity and connectivity from the physiologically most relevant early time windows remains largely unfulfilled due TMS–EEG artifacts. To circumvent the issue, many TMS–EEG studies have focused on much later (> 50 ms) events. Unfortunately, these late components are contaminated by auditory and tactile brain activations from the TMS coil (Conde et al., 2019; Siebner et al., 2019). Moreover, such long-latency responses are distant echoes from the initial earlier activations of main interest. This motivates studies on how to reduce early TMS–EEG artefacts.

Optimizing EEG recording techniques are essential in reducing the decay artifact. Previous studies have highlighted the importance of choosing electrode types (*e.g*., pellets) that minimize the TMS-induced eddy currents in them (Ilmoniemi and Kicić, 2010) or reduce the electrode–scalp impedances as much as possible (Julkunen et al., 2008), or orienting the EEG electrode leads such that induction from the TMS coil windings is minimal (Sekiguchi et al., 2011), and minimizing EEG electrode and lead movement (Veniero et al., 2009; Mutanen et al., 2013) (e.g., with plastic film wrap). Following these principles is necessary when early latencies are being studied, but this comes at the cost of long EEG preparation times. Moreover, the leads can only be oriented for one pre-determined TMS coil location/orientation per EEG preparation. However, even with the best practices, recording clean EEG signals within ∼12 ms after a TMS pulse is in many subjects not possible, resulting in loss of usable data.

Several studies have developed software means for modeling (Freche et al., 2018) and removing the decay artifact and other TMS–EEG artifacts (Mäki and Ilmoniemi, 2011; Rogasch et al., 2014; Mutanen et al., 2016; Casula et al., 2017; Wu et al., 2018; Mutanen et al., 2020; Rogasch et al., 2022; Metsomaa et al., 2024; Vergani et al., 2025). Yet, these methods do not currently offer generally applicable solutions to TMS–EEG data. Computational separation between artifacts and brain signals is still incomplete, and some correction methods may generate signals that were not in the original data (Bertazzoli et al., 2021).

In summary, the present study confirms the role of EEG lead configuration in reducing both TMS pulse and decay artifacts and shows that certain EEG lead configurations may be particularly useful.

